# Seasonal behavioral patterns of the Caquetá titi monkey (*Plecturocebus caquetensis*)

**DOI:** 10.1101/2024.01.10.573946

**Authors:** Acero-Murcia Adriana Carolina, Almario-Vaquiro Leidy, Ortega Zaida, García-Villalba Javier, López-Camacho René, Defler Thomas

## Abstract

The Caquetá titi monkey (*Plecturocebus caquetensis*) is Critically Endangered (CR) due to habitat destruction associated with extensive livestock ranching and farming. But unfortunately, information on the ecology and behavior of the species is scarce. Here we describe seasonal behavioral patterns and home range size of *P. caquetensis* in a 23-ha forest fragment in Colombia. In 2013, we collected focal animal behavioral data of a group of seven individuals of *P. caquetensis*. Overall, titi monkeys spent 33% of their time feeding, 4% locomotion, 42% resting, and 21% engaging in social interactions, such as grooming and vocalizations. However, the activity budgets differed substantially between seasons. In the dry season titi monkeys invested significantly more time feeding than in the rainy season, when they spent more time moving. They invested a similar amount of time in social interactions in both seasons. Home range size was larger during the rainy season; but the core areas of the home ranges had a similar size and they overlapped across seasons, suggesting the overall importance of this area for titi monkeys. These findings constitute the first study on the seasonal ecology and home range of this Critically Endangered species, and it identifies a priority local conservation area.

## INTRODUCTION

Studying the activity patterns of animals helps us to understand how the habitat and environment shape the time invested in different behaviors (Isbell and Young 1993; Brockman and Van Schaik 2005; Agostini *et al*. 2012). Behavioral patterns are mainly related to the availability of food resources. For example, the scarcity of resources can cause an increase in travel distances or reduce dietary breadth and time of social interactions (Hemingway and Bynum 2005). Primate studies show that environmental and ecological conditions influence dietary composition (Stevenson 2006; Nagy-R and Setz 2017), habitat use (Setz *et al*. 2013), social activity (Toro *et al*. 2020), and reproduction (Stone and Ruivo 2020; Heldstab *et al*. 2020). For example, saki monkeys (*Pithecia pithecia*) increase feeding time in the dry season, when their preferred food ítems –as immature seeds– are more abundant (Norconk 1996). In Suriname, the feeding time of bearded sakis (*Chiropotes sagulatus*) seasonally shifts due to the higher availability of seeds in the dry season (Setz *et al*. 2013). Another study shows how increasing availability of food can increase the time invested in social interactions, such as play or grooming in young howler monkeys (*Alouatta palliata Mexicana*) (Toro *et al*. 2020). The reproduction of squirrel monkeys (*Saimiri collinsi)* coincides with suitable environmental conditions and peak food availability of the Amazon Forest, in a way that increases infant survival (Stone and Ruivo 2020). Understanding feeding habits and the spatiotemporal availability of the preferred food items can help predict the impact of climate change and habitat loss on a given species. In Mesoamerican spider monkeys (*Ateles geoffroyi)* resting time increases when rainfall decreases, whereas feeding time increases with rainfall, and locomotion is negatively related to rainfall and temperature (González-Z *et al*. 2011). Furthermore, *A. geoffroyi* spends more time feeding and less time resting in forest fragments than in continuous habitats (González-Z *et al*. 2011). So, climatic conditions, habitat, and spatial and temporal distribution of food sources influence behavioral patterns and, consequently, the use of space, such as daily ranges or the home range (Milton and May 1976; De la Torre *et al*. 1995; Hanya *et al*. 2020).

Understanding how animals move and use space is key to gain an integrative comprehension of how they behave (Nathan *et al*. 2008). Basic aspects of describing the home range include the space where an individual conducts daily activity, excluding occasional explanatory incursions into other areas within the home range; the core area is the area used most intensively (Burt 1943; Fryxell *et al*. 2014). Primates often change the time invested in locomotion according to the season. During the lean season individuals increase the time moving or searching in resource-rich areas, thus enlarging their home range area (Brockman and Van Schaik 2005). Home range size appears to be more flexible among New World primates, because they increase their dietary diversity in response to food availability, contrary to primate communities from Africa, Asia, or Madagascar (Brockman and Van Schaik 2005). For titi monkeys, as for many other social mammals that live in groups, home range is estimated at the group level (Milton and May 1976; Reyna-H and Chapman 2019). In Japanese macaques (*Macaca fuscata*) home range size is affected by food availability because these primates spend more time searching for high-energy items like flowers and fungi, and this directly lengthens the duration of locomotion and home range size during the winter (Hanya *et al*. 2020). In contrast, the home range of *Leontocebus nigricollis graellsi* in Ecuador becomes approximately 25% smaller in the rainy season, probably due to the distribution and availability of fruits in this area (De la Torre *et al*. 1995). This same pattern is also observed in *Cheracebus torquatus* (Hoffmannsegg, 1807) and *Cebus albifrons* (Humboldt, 1812) in Panama and Peru, where changes in the home range size result from a higher productivity of resources like fruits and insects during the rainy season (De la Torre *et al*. 1995). Identifying core areas is pivotal to understanding why home ranges are more extensively utilized (Asensio *et al*. 2012a; Fleagle 2013; Fryxell *et al*. 2014). For example, *Cheracebus lugens* has a core area of 8.75 ha, where most feeding bouts take place, accompanied by duet vocalizations, probably to defend the fruit resource of *Virola michelii* Heckel (Palacios *et al*. 1997). In addition, spider monkeys (*Ateles geoffroyi*) in Costa Rica have five core areas of approximately 9.4 ha that correspond to sites with higher habitat quality than non-core areas (Asensio *et al*. 2012b). In general, previous studies provide evidence that core areas have key resources, like mature forests that provide a diversity of food trees and sleeping sites for the establishment of a particular group (Asensio *et al*. 2012b).

In Colombia, studies on behavioral patterns of titi monkeys (*Plecturocebus, Cheracebus*) have focused on the departments of Meta for *Plecturocebus cupreus* (Spix, 1823) (Mason 1965, 1971; Polanco *et al*. 1994; Porras 2000), *Plecturocebus ornatus* (Mason 1968), and *Plecturocebus moloch* (Mason 1966; Moynihan 1966); in the department of Vichada for *Cheracebus lugens* (Defler 1983; Defler and Carretero-Pinzón 2018); on Vaupés for *C. lugens* (Palacios *et al*. 1997; Palacios and Rodríguez 2013; Defler and Carretero-P 2018); and the Amazonas for *C. lucifer* (Defler and Carretero-P 2018). Primates of this subfamily are characterized as highly territorial (Moynihan 1976; Robinson 1979; Mitani and Rodman 1979) and pair-bonded (Mitani and Rodman 1979; Bicca-M and Heymann 2013; Fernandez-D *et al*. 2020), displaying characteristic vocal duets and choruses (Moynihan 1966; Adret *et al*. 2018). The groups are usually composed of one adult pair and their offspring, usually not exceeding five (Bicca-M and Heymann 2013) or seven individuals (Acero-M *et al*. 2018).

The natural history of the Caquetá titi monkey (*Plecturocebus caquetensis*) is poorly known (Acero-M *et al*. 2018; Acero-M *et al*. 2021), even though it is highly threatened by extensive livestock farming, illicit crops, and illegal logging, which deforested 29,800 ha in Caquetá department in 2019 (García and Defler 2013; Defler *et al*. 2016; Murad and Pearse 2018; Castro-N *et al*. 2019). Our aim in this article was to evaluate the seasonal patterns of behavior and home range size of *Plecturocebus caquetensis*. Given the behavior of other species of primates, including congeneric species, we expected *P. caquetensis* to show seasonal behavioral variations, with a tendency to reduce the time invested in locomotion and increase the time in feeding during the dry season (Kinzey and Norconk 1993). Thus, we expected that *P. caquetensis* minimizes energetically costly activities in the dry season, following an energy-minimization strategy (Agostini *et al*. 2012; Nagy-R and Setz 2017; Del Toro *et al*. 2019; Souza-A *et al*. 2021). As a consequence of increasing locomotion, we also expected a more extensive home range during the rainy season.

## METHODS

### Study site

This study was conducted in 2013 in a 23-ha secondary forest fragment in Playa Rica, Municipality of Valparaíso-Caquetá, Colombia (01°04’ North, 75°36’ West, 224 m a.s.l.; Fig. 1a). The study site was a mosaic of forest fragments within a matrix of cattle pasture and *Hevea brasiliensis* monoculture. The Fabaceae, Moraceae, and Myristicaceae families were the most abundant in the study forest. Maximum canopy height was 40 m (Acero-Murcia *et al*. in prep.). The climate was tropical humid, with a maximum annual average temperature of 26.2 °C, and a mean annual rainfall of 260 mm (Fig. 2, IDEAM 2014). The rainfall pattern was unimodal, with a maximum from April to July and a minimum in the second semester of the year (Guzmán *et al*. 2014; Urrea *et al*. 2019).

**Figure 1.**
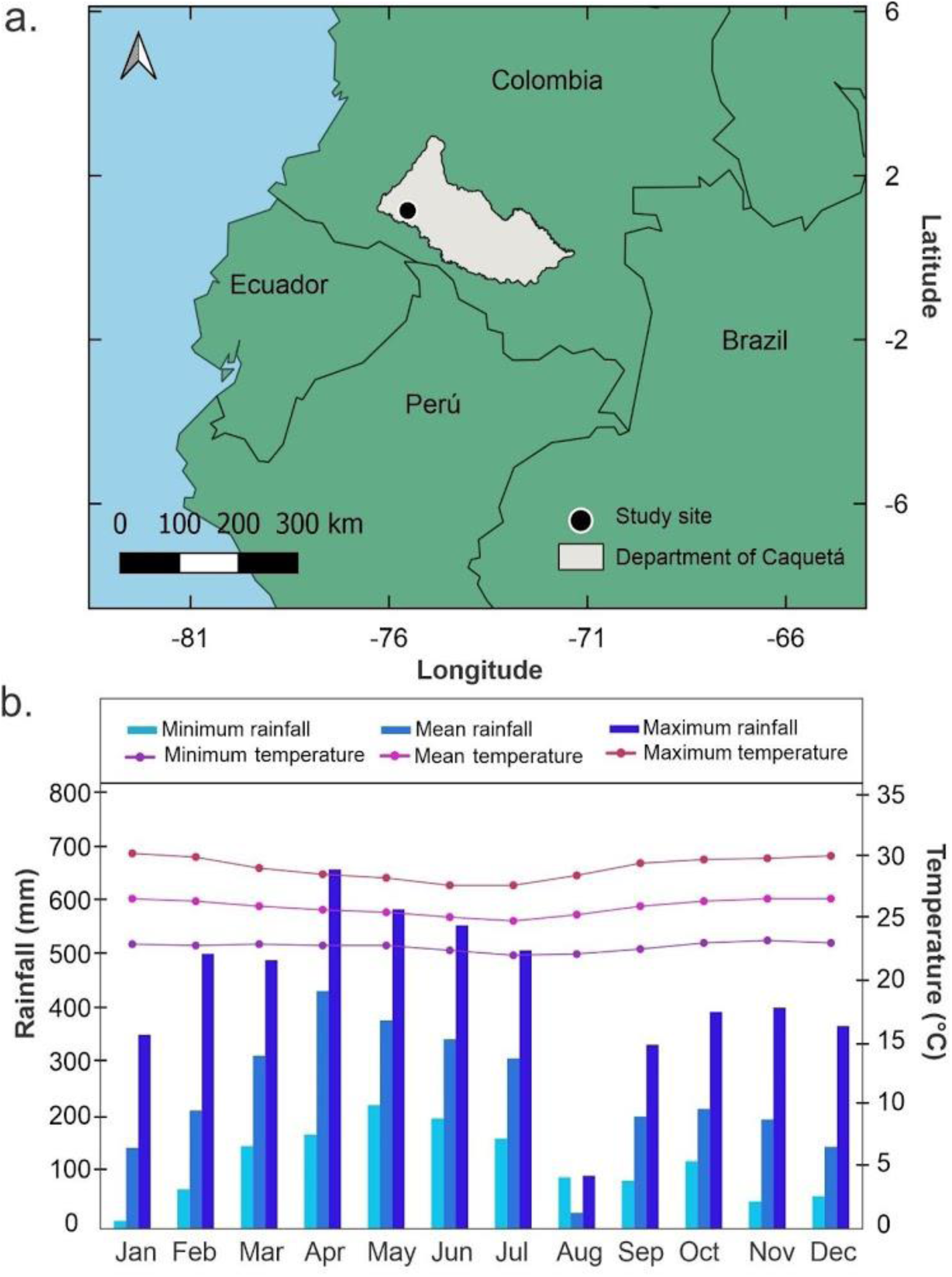
(**a**). Study site in Playa Rica, municipality of Valparaiso-Caquetá, Colombia. (**b**). Rainfall (bars) and temperature (dots) values from Valparaiso station (IDEAM 2014).

**Figure 2.**
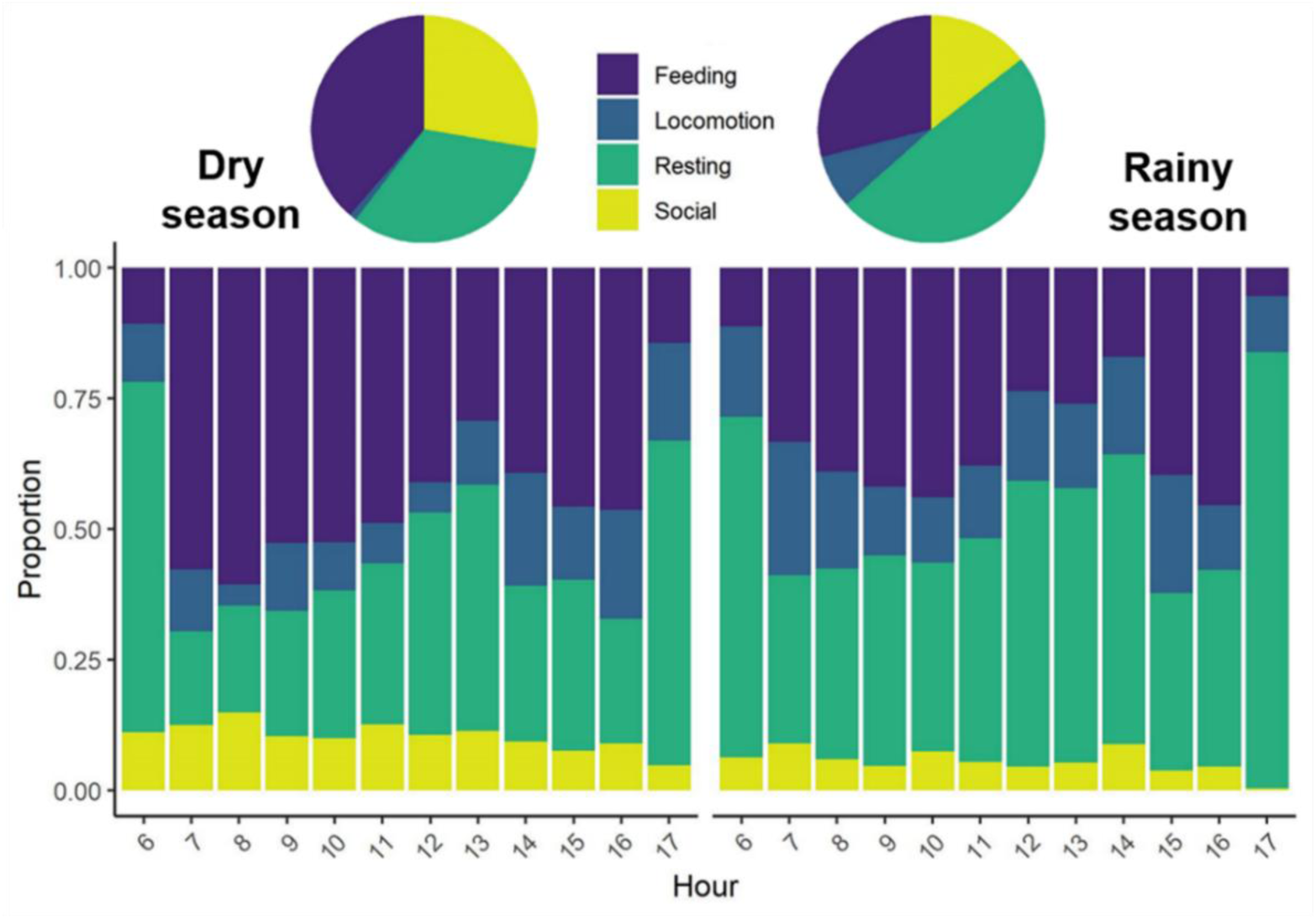
The bar chart shows the relative proportion of time that the *Plecturocebus caquetensis* group spent on each behavior throughout the day during the dry and rainy seasons. The pie charts represent the total percentage of time spent on each behavior by season.

### Study group and monitoring

We monitored a group of Caquetá titi monkeys (*P. caquetensis*) from February to December 2013. During the study, the composition of the group varied from six to seven individuals. The group was composed of one adult female, one adult male, one sub-adult male, two juveniles (male and female), one infant, and a newborn (born in early October 2013). The observations did not focus on the infant or the newborn. In addition, we did not discriminate by sex in the focal individuals. We collected data from 06:00 – 18:00 h. In total we observed the group for 47 days, totaling 557.54 hours.

We registered four behavioral categories continuously during ten minutes for each focal animal sample (Altmann 1974), interspersed with five-minute lapses used to choose another individual or to register data, thus obtaining four independent focal samples per hour for different individuals (Table 1). We shifted the focal individuals for each sampling register. When we lost a focal individual from sight during a focal sampling, another individual of the group was chosen to begin a new focal sample. The order of focal animals was determined randomly with at least four sampling registers left between separate observations of the same individual.

**Table 1.** Ethogram of recorded behaviors for *Plecturocebus caquetensis* in Caquetá, Colombia.

| Behavioral category | Description |
| --- | --- |
| **Feeding** | The primate is searching for, manipulating, or ingesting fruits, leaves or insects on a single tree, several trees, or sometimes on the ground looking for insects or other arthropods. Licking water directly from the leaves of some epiphytes by manipulating the leaf with both hands as a funnel near the mouth. |
| **Locomotion** | The primate moves, regardless of whether it is in vertical or horizontal displacement. It is usually a quadruped movement, occasionally jumping from branch to branch or climbing vertically. |
| **Resting** | The primate shows no movement, usually leaning on its front and rear limbs on a branch, with the back semi-straight as if sitting (Moynihan 1966, Mason 1966), or with the ventral part of the body on a horizontal branch, usually with the lower extremities hanging down and the upper extremities on the branch (Moynihan 1966). |
| **Social interactions** | This category includes vocalization, play, parental care, breastfeeding, reproductive behavior, rubbing the body and grooming. It includes interactions involving one (i.e., allogrooming, vocalization) or more individuals (i.e., parental care). |

To facilitate habituation and monitoring of the monkeys, we opened trails separated by 25 m and intersecting at right angles throughout the study area. This grid facilitated the localization of monkeys in the area, allowing us to locate key grids for food or rest trees. To estimate the home range and core area for both seasons (See statistical analysis), we recorded geographical coordinates every 30 minutes using a GPS device (Garmin Oregon 400t), starting at 06:00 h and continuing throughout the day.

### Ethical note

This study was carried out within the project “Conservation Plan of the Mico Bonito del Caquetá *Callicebus caquetensis* (Defler, Bueno and Garcia 2010)”, which follows the ethical standards of Colombian law for working with primates and the American Society of Primatologists Principles for the Ethical Treatment of Non-Human Primates (Setchell and Curtis 2003; Buchanan *et al*. 2012).

### Data analysis

According to the unimodal climate regime (Fig. 1b), the sampling period was divided into: (1) dry season, totaling six months (February, August–December) and (2) rainy season with five months (March–July); January was not included because there were no data. Monthly precipitation values (mm) and temperature (°C) values were obtained from Valparaíso-Caquetá Meteorological Station (IDEAM 2014).

To estimate the daily path and length of the time budgets for the different behaviors and subsequently to assess the seasonal influence, we calculated the relative frequency of each behavior at each hour of the day (06:00 – 18:00 h) for each season. The relative frequency for each category was estimated as the proportion of time in this category divided by the total time registered in the correspondent hour and season (dry or rainy). Thus, for each season, the contribution of each behavioral category at each hour was estimated as the time that the animal spent in this category divided by the total time registered for a particular hour and season.

To analyze the effect of the season (rainy vs dry) on the daily patterns of the time budgets of the four behavioral categories, we modeled the proportion of time spent in each behavior using a Dirichlet regression (Douma and Weedon 2018). Data of proportions have the particularity of being constrained to an interval (0-1 in the proportions, or 0-100 in the percentages), which results in their variances not being constant across the values of the predictor (Douma and Weedon 2018). Transformations are often performed to overcome these limitations and to be able to fit linear models, but they can provide biased estimates of the coefficients (Douma and Weedon 2018). Thus, the Dirichlet regression is the most suitable available method to model data on proportions with more than two categories, as in our case, with four behavioral categories (feeding, resting, locomotion, and social interactions). This method has been previously used to estimate time budgets of seabirds for four activities (Regular *et al*. 2014). A detailed guide to conduct Dirichlet regression is available in Douma and Weedon (2018). We conducted the Dirichlet regression using the DirichletReg package (Maier 2014), fitting a model on the proportion of time for each activity (response variable) and the season and hour of the day (scaled) as moderators. We used the “alternative” parametrization, where the vector of expected values (μ) is modelled as a function of covariates and a precision parameter (ϕ) (Douma and Weedon 2018). We fitted different models, starting with the most complex (including the interaction between season and hour, both in the expected values and in the precision parameter) and simplified the model in a backward stepwise approach, based on the variation of AIC (Akaike Information Criterion) between models (contrasted via ANOVA). The most parsimonious model in terms of AIC resulted as follows: proportion ∼ season + hour, where the precision parameter was constant.

Finally, we estimated the home range of the group for each season using two estimators, the 95 % minimum convex polygon (MCP) (Mohr 1947) and the kernel density estimator (KDE), using the 95 % and 50 % (to estimate the core area of the home range) isopleths of probability, the reference smoothing parameter (href) and calculating the diffusion parameter by maximum likelihood (Worton 1989). We obtained the home range estimation using the adehabitatHR package (Calenge 2011). All analyses were performed in the R environment (R Core Team 2019).

## RESULTS

In general, the group invested more time resting and feeding, followed by social interactions and with a lower temporal investment in locomotion. The activity budget for the dry and rainy season in terms of the observed behavioral categories for *Plecturocebus caquetensis* is reported in Table 2.

**Table 2.** Activity budget of Caquetá’s titi monkey, *Plecturocebus caquetensis*, on each of the four studied behavioral categories. Mean and ± Standard deviation percentages of the time spent by category.

| Behavior | Dry season | Rainy season | Total |
| --- | --- | --- | --- |
| **Feeding** | 38.58 ± 13.14 | 28.96 ± 10.98 | 33.07 ± 12.81 |
| **Locomotion** | 1.05 ± 0.42 | 7.60 ± 3.06 | 4.16 ± 3.83 |
| **Resting** | 32.66 ± 15.77 | 48.97 ± 16.04 | 41.84 ± 17.81 |
| **Social Interactions** | 27.71 ± 5.09 | 14.47 ± 5.02 | 20.93 ± 8.53 |

The model including the main effects of season and hour and a constant precision parameter fitted the data with the same quality as the more complex models, so we kept this model among the other candidates (Table 3). The season significantly influenced the time that the group invested in locomotion, resting, and feeding; in addition, behaviors such as resting and feeding changed with the hour (Table 4).

**Table 3.** Candidate Dirichlet’s regression models for fitting the proportion of time invested on the four studied behaviors (feeding, locomotion, resting, and social interactions). ANOVA (of the deviances) were conducted to compare each model with the simplest one.

| Model | AIC | Deviance | p-value |
| --- | --- | --- | --- |
| proportion ~ season + hour (dispersion term constant) | -189.9 | -209.92 | - |
| proportion ~ season + hour (dispersion term = season) | -186.9 | -210.89 | 0.615 |
| proportion ~ season + hour (dispersion term = hour) | -188.9 | -210.86 | 0.332 |
| proportion ~ season + hour (dispersion term = season + hour) | -186.9 | -210.89 | 0.615 |
| proportion ~ season + hour (dispersion term = season + hour + season x hour) | -191.1 | -217.13 | 0.066 |
| proportion ~ season + hour + season x hour (dispersion term constant) | -185.1 | -211.10 | 0.759 |
| proportion ~ season + hour + season x hour (dispersion term = season) | -183.1 | -211.12 | 0.879 |
| proportion ~ season + hour + season x hour (dispersion term = hour) | -183.8 | -211.77 | 0.763 |
| proportion ~ season + hour + season x hour (dispersion term = season + hour) | -181.8 | -211.80 | 0.866 |
| proportion ~ season + hour + season x hour (dispersion term = season + hour + season x hour) | -186.4 | -218.41 | 0.204 |

**Table 4.** Results of the Dirichlet regression model (alternative parametrization) on the proportion of time spent by *Plecturocebus caquetensis* according to the season and the hour of the day. Coefficients for feeding are not shown because it is the reference category. Log-likelihood: 105 on 10 df, AIC: –189.9, BIC: –178.1.

| Variable | Coefficient | Estimate | SE | Z | p-value |
| --- | --- | --- | --- | --- | --- |
| **Locomotion** | Intercept | -2.595 | 0.284 | -9.134 | < **0.001** |
|  | Season (rainy) | 1.384 | 0.352 | 3.927 | < **0.001** |
|  | Hour | -0.038 | 0.159 | -0.237 | 0.813 |
| **Resting** | Intercept | -0.131 | 0.146 | -0.899 | 0.369 |
|  | Season (rainy) | 0.742 | 0.203 | 3.650 | < **0.001** |
|  | Hour | 0.214 | 0.104 | 2.056 | **0.039** |
| **Social interactions** | Intercept | -0.233 | 0.150 | -1.555 | 0.119 |
|  | Season (rainy) | -0.421 | 0.240 | -1.759 | 0.079 |
|  | Hour | -0.175 | 0.113 | -1.549 | 0.121 |
| **Precision model** | Intercept | 3.088 | 0.167 | 18.51 | < **0.001** |

*Plecturocebus caquetensis* spent a significantly higher proportion of time moving (locomotion) and resting during the rainy season than in the dry season, when they invested proportionally less time feeding (Table 4; Fig. 2). Resting behavior changed throughout each hour and for each season. However, the daily patterns were constant between seasons; that is, they had rest peaks at dawn, at noon, and at dusk (Fig. 2). Social interactions did not differ between seasons and hours. Although there was evidence of an increase in this activity during the dry season, it was not statistically significant (Table 4; Fig. 2). Finally, home range estimates for *P. caquetensis* were larger during the rainy (6.67 ha) season than during the dry (4.16 ha) season (Fig. 3).

**Figure 3.**
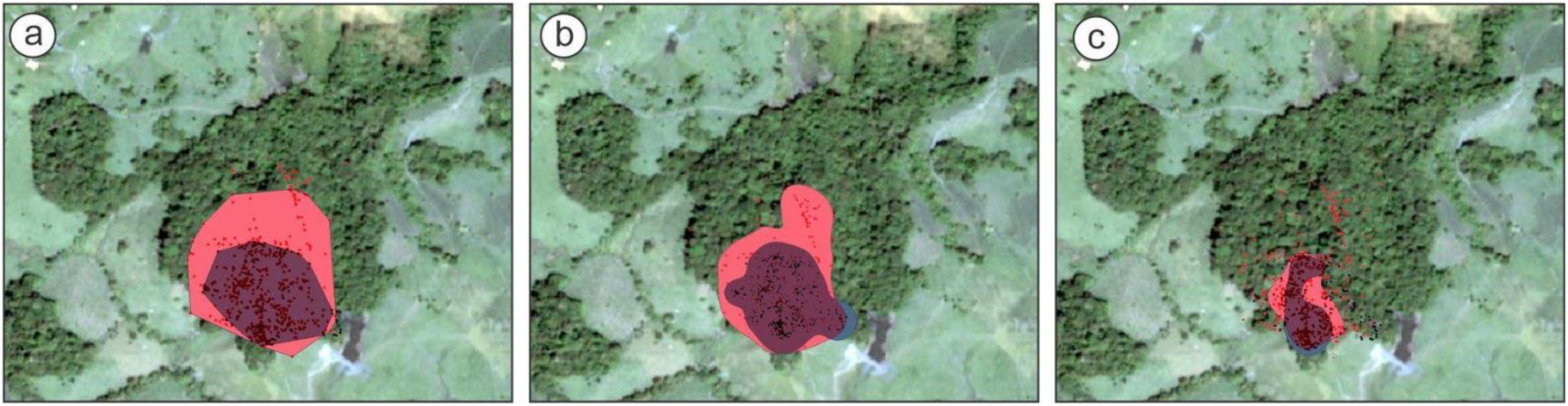
Home range for *Plecturocebus caquetensis* in the rainy (red) and the dry (gray) seasons. (**a**) Estimation of MCP 95% (2.74 ha dry season, 5.58 ha rainy season), (**b**) estimation of KDE 95 % (4.16 ha dry season, 6.77 ha rainy season), and (**c**) estimation of KDE 50% (1.13 ha dry season, 1.65 ha rainy season).

## DISCUSSION

Our findings showed that the general behavior of *P. caquetensis* were quite similar to those reported for the Amazon region and for the genus *Callicebus sensu lato*. For other species the main observed activities are resting, and feeding for the following species: *C. personatus* (Kinzey and Becker 1983); *C. torquatus* (Easley 1982; Palacios *et al*. 1997; *P. cupreus* (Müller 1995; Kulp and Heymann 2015). For seasonal behavioral patterns, *P. caquetensis* spent more time feeding and less time moving during the dry season and, thus, the increased time for locomotion produced a home range size that was greater in the rainy season, which corroborates our hypotheses.

### Feeding and Locomotion

*Plecturocebus caquetensis* spent more time feeding during the dry season than in the rainy season, probably because of the increased availability of young leaves (*Sorocea muriculata, Davilla* sp*, Tapirira* sp.), seeds (*Pourouma bicolor*, *Siparuna* sp.), fleshy fruits (*Bellucia pentamera*), exocarps (*Socratea exorrhiza*), flowers (*Eschweilera punctata*), and arthropods (Acero-M *et al*. 2018). Among other trophic resources, *Plecturocebus* actively selects for seeds (i.e., uses this resource in a higher frequency than available) (Kinzey and Becker 1983; Palacios *et al*. 1997; Alvarez and Heymann 2012; Palacios and Rodriguez 2013; Acero-M *et al*. 2018). This leads *P. caquetensis* to increase the time invested in feeding during the dry season. For instance, in the Atlantic Forest of Brazil, the diet of one group of the black-fronted titi monkey *Callicebus nigrifrons* is dominated (90%) by seeds during the dry season, while in other groups at the same study site and season, seeds contribute 72.8%, 46.7% and 41.5% of the diet (Dos Santos *et al*. 2012). In addition, Caselli and Setz (2011) also report an increase in feeding time in the dry season also for *C. nigrifrons*. Similarly, the Rio Mayo titi (*P. oenanthe*) in the Peruvian Andes shows a high consumption of insects during the dry season (DeLuycker 2012). So, titi monkeys reduce time invested in locomotion when fleshy fruits are scarce, but they compensate for this demand by eating lipid-rich seeds as an alternative high-nutritional resource (Lambert 1998) widely available during this season (Kinzey and Norconk 1993). Time processing seeds is higher compared to other food ítems, which can lead to reduce locomotion as a strategy for maximizing energy balance (Heiduck 2013). For example, *C. nigrifrons* in the Atlantic Forest travels less in the dry season but increases its diet breadth by consuming seeds and leaves. This can be considered as a strategy to minimize energy expenditure (Nagy-R and Setz 2017). Moreover, the *P. caquetensis* group increased locomotion during the rainy season, probably trying to find resources like ripe fruits that are rich in sugars and are more easily digested than leaves (Strier 2007). These results agree with those for *C. coimbrai* that had a greater number of locomotion records due to the low supply of resources like fleshy fruits in the rainy season (Souza-A *et al*. 2011). Likewise, the black-tufted marmoset (*Callithrix penicillata*) invests more time in locomotion during the rainy season (Vilela and Faria 2004), and *C. flaviceps* travels for longer periods in the rainy season when fruits and flowers are scattered (Ferrari 1988). In general, the investment of time for locomotion varies depending on the quality of the habitat, the availability of resources, and the distribution of optimal sleeping sites (Kinzey and Becker 1983; Heiduck 2013).

### Resting

The proportion of time that *P. caquetensis* invested in rest was higher at dawn, at noon, and at dusk, when individuals reduce the time in other behaviors. This diurnal pattern is similar for most primates (Fleagle 2013). In relation to the season and hour, we found differences in times invested for each behavior, with resting behavior increasing in the rainy season and changing hourly. For example, Coimbra-Filho’s titi monkey spends more time resting in the rainy than in the dry season in a forest fragment. This is probably due to a strategy for minimizing energy expenditure when ripe fruits are scarce (Souza-A *et al*. 2021), since resting requires less energy than other behaviors (Heiduck 2013). In primates, diet composition, temperature, and fragment size influence resting time, principally because these variables modify the time needed for digestion (Korstjens *et al*. 2010). In fact, the Geoffroyi’s spider monkeys (*Ateles geoffroyi*) invest more time resting during the dry season, when the consumption of young leaves is known to have an effect on the time of digestion (Chaves *et al*. 2011). During resting peaks, some species, such as *Cheracebus torquatus*, increase grooming at the sleeping site (Kinzey *et al*. 1977; Kinzey and Wright 1982). In general, our results for *P. caquetensis* agree with the behaviors reported for other Callicebinae species, which spend around 29% to 63% of their time resting (Bicca-M and Heymann 2013).

### Social Interactions

The time invested in social interactions for *P. caquetensis* differed between seasons. There are two main hypotheses to explain the seasonality of social interactions in primates. The first one predicts that nutritional stress associated with scarcity of resources may lead to a reduction in social behavior, since greater competition for food will lead to disperse the distribution of individuals within groups, therefore reducing the time invested in behaviors such as grooming or other social activities during the dry season (Foley 1987). For example, howler monkeys in Mexico invest a significantly greater amount of time in social interaction during the season of abundance rather than in the lean season, and costly activities like play are more frequent in the young members (Agostini *et al*. 2012). Probably, high energy-demanding behaviors can be carried out during the season of greater abundance, when consumption of high-quality food like fruits is more frequent than in the lean season, and this allows social activities to increase (Alberts *et al*. 2005; Agostini *et al*. 2012). In *P. caquetensis,* social activities like play were observed on different occasions; and, as expected, there was greater participation of young individuals (*pers.obs*). However, this did not represent increased time when compared with grooming. Another social activity that primates frequently show is grooming, which reinforces social bonds and is characteristic in *Callicebinae* (Jones and Anderson 1978; Kinzey and Wright 1982). In *P. caquetensis*, all group members showed this behavior. Grooming in *P. caquetensis* illustrated a trend that increased during the dry season compared to the rainy season, although the difference was not statistically significant. Nonetheless, this result showed that the time invested in social interaction increased in the dry season. This could be explained because grooming does not require such a high energy expenditure as does playing; and because primates can reduce resting time in order to maintain social activities (Dunbar 1992) like grooming in the lean season, as we found here.

During the dry season the parents of our study group gave birth to a newborn. This event increased care activities, including breastfeeding and vigilance towards other coexistent groups, such as saki monkeys (*Pithecia milleri*) (*pers. obs*). A second hypothesis for explaining our results is that time budgets can adapt to changes in resource availability, predicting that less time will be used for resting than for social behavior to accommodate the increased time demanded for foraging during the dry season. This is known as the ‘social glue’ hypothesis (Dunbar and Dunbar 1988; Dunbar 1992). For example, baboons (*Papio cynocephalus* Linnaeus, 1766) keep their social time and reduce resting time to accommodate increased foraging demands during the lean season; but they sacrifice social activities over the time in lower-quality habitats that demand more investment for foraging (Alberts *et al*. 2005). This supports the idea that time invested in each season can vary according to habitat quality. If this is the case, we expect that *P. caquetensis* populations will be affected in the long term by deforestation and the loss of food sources, leading to an increase of time for searching and a decrease of time for social activities.

### Home range and core areas

The home range size of *P. caquetensis* (2-7 ha) was similar to that reported for other titi monkeys such as *Plecturocebus cupreus* (3.29 – 4.18 ha, Robinson 1979), *P. discolor* (I. Geoffroy Saint-Hilaire and Deville 1848) (3.3 ha, Carrillo-Bilbao *et al*. 2005), *P. oenanthe* (2 ha, DeLuycker 2007)*, P. ornatus* (0.3 – 0.5 ha, Mason 1966, 1968), *P. brunneus* (Wagner 1842) (<2 ha and 3 ha, Lawrence 2003, 2007), or *P. olallae* (Lönnberg 1939) (7.2 ha, Martinez *et al*. 2015) in fragmented or degraded forest patches. However, other studies conducted in continuous forests report home range sizes larger than 10 ha (11.4 ha, Kulp and Heymann 2005; 22 ha, Palacios and Rodriguez 1997; 22 ha, Heiduck 2002). In terms of seasonality, the study group travelled more, thus increasing its home range, in the rainy than in the dry season. Studies have demonstrated that primates can adjust the activity budget and the home range size in fragmented forests in relation to food availability. For instance, the golden-faced Saki (*Pithecia chrysocephala*) travels more in the rainy season when the availability of fleshy fruits is higher (Setz 1993). Also, *Callithrix flaviceps* shows larger home ranges in the rainy season due to the availability of prey insects, contrary to *Callithrix aurita*, that shows a different pattern, supporting that home range size may also vary due to other environmental situations affecting habitat quality (Ferrari *et al*. 1996). Another example is *Leontocebus nigricollis graellsi* who reduces its home range size during the dry season in flooded forests and increases its home range size during the rainy season in non-flooded ones (De la Torre *et al*. 1995). Home range reduction in the dry season can be explained because there is higher productivity of resources such as fruits and insects in the rainy season providing enough resources in a small area. This was also reported for *Cheracebus torquatus* and *Cebus albifrons* (De la Torre *et al*. 1995). On the other hand, the core area of *P. caquetensis* was similar in size and location between the rainy and the dry season. This area included sleeping sites and key feeding trees for *P. caquetensis* (*pers. obs*). A study on *P. discolor* in Yasuni (Ecuador) shows home range (1.7 – 6.4 ha) and core area (0.8 – 3.3 ha) sizes similar to our results (Van Belle *et al*. 2020). In addition, a study on the influence of resource distribution and ranging patterns in *Aotus azarae* (Fernandez-D 2016) suggests that during the lean season or in bottleneck conditions, the monkeys have access to similar resources irrespective to the size of their home ranges or core areas, because these areas offer the minimum nutritional requirements in the dry season.

### Conclusion and perspectives in conservation

*Plecturocebus caquetensis* can be considered an energy maximizing species since the individuals reduce costly behaviors (like locomotion) and increase other activities (like feeding) in the dry season. The activity budget and home range size follow the pattern of adaptability in primates that inhabit fragmented or isolated forests. To minimize the effect of habitat loss for the Caquetá titi monkey (*P. caquetensis*), protected areas should be established because the department of Caquetá had a deforestation rate of 0.77% between 2000 and 2020, about twice as high as estimated from previous studies for the other South America tropical regions (Murad and Pearse 2018). Approximately 13.8% of pristine forests have been lost, resulting in *P. caquetensis* being one of the most threatened species due to habitat loss in Colombia (Boyle 2014; Defler *et al*. 2017; Murad and Pearse 2018). Our data are limited, and further assessments on the effect of forest loss and fragmentation on the species’ populations are needed. Thus, we encourage research on the dispersal capacity of *P. caquetensis* in fragmented forests inserted in pasture matrices for livestock, and monitoring of its populations throughout the species distribution range.

## CONTRIBUTION OF THE AUTHORS

ACAM, TRD, JG, LAV and RL conceived and designed the study, ACAM and LAV collected the data, ACAM and ZO conducted the analyses, and ACAM, LAV, ZO wrote the manuscript with input from the other authors.

## ACKNOWLEDGEMENTS

We thank Marta Bueno, Paul Bloor, Edwin Trujillo and Carolina Ibañez for their contributions to the Conservation Plan. Also, we thank the institutions Fundación Herencia Natural, Universidad Nacional de Colombia, Universidad Distrital Francisco José de Caldas, Instituto de Ciencias Naturales de la Universidad Nacional de Colombia and Universidad de la Amazonia for the infrastructure and partnership.

## FUNDING

This work was supported by Pacific Rubiales Energy S.A. Foundation (Conservation Plan of the Mico Bonito del Caquetá “*Callicebus caquetensis*”). ACAM and ZOD, were supported by the Coordenação de Aperfeiçoamento de Pessoal de Nível Superior – Brazil (CAPES) for the doctoral (Financing code 001) and postdoctoral scholarships PNPD/CAPES (process 1694744).

## CONFLICT OF INTERESTS

The authors declare that there is no conflict of interests.

## DATA AVAILABILITY STATEMENT

All data generated, and scripts for this study are available in Data Dryad: https://doi.org/10.5061/dryad.1jwstqjt9

## Notes

### Competing Interest Statement

The authors have declared no competing interest.

